# A highly efficient human cell-free translation system

**DOI:** 10.1101/2023.02.09.527910

**Authors:** Nikolay A. Aleksashin, Stacey Tsai-Lan Chang, Jamie H. D. Cate

## Abstract

Cell-free protein synthesis (CFPS) systems enable easy *in vitro* expression of proteins with many scientific, industrial, and therapeutic applications. Here we present an optimized, highly efficient human cell-free translation system that bypasses many limitations of currently used *in vitro* systems. This CFPS system is based on extracts from human HEK293T cells engineered to endogenously express GADD34 and K3L proteins, which suppress phosphorylation of translation initiation factor eIF2α. Overexpression of GADD34 and K3L proteins in human cells significantly simplifies cell lysate preparation. The new CFPS system improves the translation of 5’ cap-dependent mRNAs as well as those that use internal ribosome entry site (IRES) mediated translation initiation. We find that expression of the GADD34 and K3L accessory proteins before cell lysis maintains low levels of phosphorylation of eIF2α in the extracts. During *in vitro* translation reactions, eIF2α phosphorylation increases moderately in a GCN2-dependent fashion that can be inhibited by GCN2 kinase inhibitors. We also find evidence for activation of regulatory pathways related to eukaryotic elongation factor 2 (eEF2) phosphorylation and ribosome quality control in the extracts. This new CFPS system should be useful for exploring human translation mechanisms in more physiological conditions outside the cell.

## Introduction

Cell-free protein synthesis (CFPS) systems have become increasingly important to advance biotechnological and fundamental research (*1*). In CFPS systems that use cell-free extracts, an mRNA of interest can be translated in conditions that recapitulate protein biosynthesis *in vivo* and reveal insights into the translation process. Cell extract-based translation systems can also be used to overcome the inherent limitations of cell-based experiments by removing cellular membrane and natural homeostasis mechanisms that prevent the screening of many translational parameters. Whereas cellular growth requires defined conditions, cellular extracts may tolerate substantial genetic or proteomic manipulation and unnatural or toxic components (*2*). In addition to functional and structural studies (*3*), proteins expressed in cell-free extracts can be used for selective and site-specific labeling, stabilization of membrane proteins in a soluble state, and for optimizing the production of toxic proteins (*4*). *In vitro* translation systems also allow high-throughput screening of thousands of individual proteins translated from mRNA libraries (*5, 6*).

To enable high levels of protein synthesis, CFPS systems require supplementation with exogenous amino acids, cofactors (proteins, small molecules, and inorganic ions), and energy resources (*1*), in addition to the translation templates provided in the form of mRNA, or DNA in the case of transcription-translation coupled systems. Many organisms have been used successfully as the source for extracts to prepare *in vitro* translation systems. *Escherichia coli* and wheat germ extracts have the highest protein production yields among the cell-free translation systems and are widely used to produce recombinant proteins (*1*). However, these systems often do not recapitulate conditions needed to translate mammalian proteins of interest.

To date, mammalian extracts have been limited to human cell lines and rabbit reticulocyte lysates (RRL). While these mammalian systems may be an adequate model for fundamental translation studies, and can provide a native environment for protein folding and post-translational modification, several shortcomings limit their functionality (*1*). Preparing RRL extracts requires labor-intensive maintenance, treatment, and sacrifice of animals, while commercially available lysates are expensive (*7*). Using rabbits as a source also limits the possibilities for genetic manipulations, including the ability to knock out or enrich specific proteins. Furthermore, RRL derives from a highly differentiated and specialized animal tissue, which limits the scope of its translation regulation mechanisms. Finally, RRL endogenously contains a very high concentration of globin mRNA, which outcompetes translation of exogenously added templates unless the RRL is pretreated with a nuclease (*7*). By contrast, the use of human cultured cells provides more flexibility, by allowing for precise genetic manipulation and the use of many different cell types (*8–10*). However, both RRL and human cell line-derived extracts have relatively low protein yields, which limits their functionality. Therefore, highly efficient human-based translation systems are needed to overcome the limitations of presently used mammalian CFPS systems.

A common limitation of presently-available mammalian translation extracts is the attenuation of translation initiation due to the phosphorylation of translation initiation factor eIF2 on subunit eIF2α. During translation initiation, eIF2 delivers initiator tRNA to the 40S ribosomal subunit in a GTP-dependent manner. After mRNA start codon recognition, eIF2-GDP is released from the ribosome and is subsequently converted to eIF2-GTP by the guanine nucleotide exchange factor eIF2B. The phosphorylation of eIF2α, which generally occurs in cells during stress (*11–13*), increases the affinity of eIF2 for eIF2B by nearly 100-fold, resulting in the sequestration of eIF2 from the translating pool and inhibition of translation (*14*). In mammalian cells, four known eIF2α-specific kinases can phosphorylate serine 51 of eIF2α (Fig. 1A). The PKR kinase is activated by double-stranded RNA, as found during viral infection or during *in vitro* transcription (*15–18*). During stress, eIF2α may also be phosphorylated by PKR-like endoplasmic reticulum kinase (PERK), general control non-derepressible-2 (GCN2) kinase, and heme-regulated HRI kinase (*14*) (Fig. 1A). Phosphorylation of eIF2α can be bypassed in human cell extracts by the addition of the human GADD34 protein, an eIF2α-specific adapter for PP1 phosphatase, and/or by vaccinia virus K3L protein, a substrate for eIF2α-specific kinases that can act as a decoy (*19–22*) (Fig. 1A). While eIF2α phosphorylation is thought to be the main limiting factor in CFPS systems, attenuation of the activity of other translation factors has not been studied in depth.

**Figure 1.**
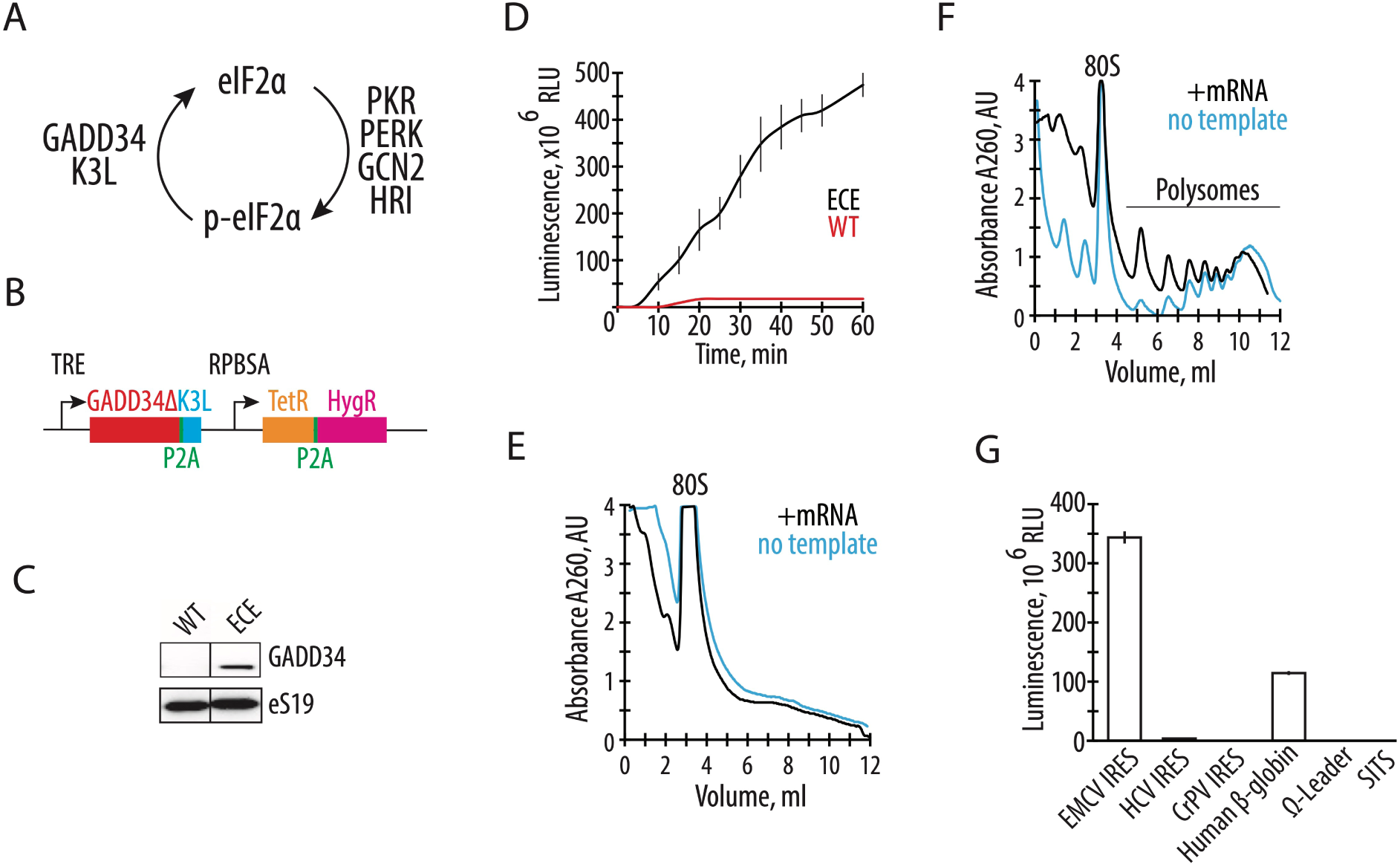
Endogenously expressed GADD34Δ and K3L increase *in vitro* translation activity of the human cell extract. **A,** Schematic of the role of GADD34 and K3L in counteracting eIF2α phosphorylation by eIF2α kinases. **B**, Diagram of the sleeping beauty-based construct used for expression of GADD34Δ (which lacks the N-terminal 240 amino acids) and K3L, which was integrated into the genome of HEK293T cells. TRE denotes a tetracycline (or doxycycline) responsive promoter that controls the expression of GADD34Δ and K3L, separated by the P2A sequence. The synthetic constitutive promoter RPBSA drives the expression of the fusion construct of tet repressor together with the hygromycin resistance gene, separated by the P2A sequence. **C,** Western blot showing expression of GADD34Δ in the engineered cell extract (ECE) but not in the WT HEK293T cells, with ribosomal protein eS19 serving as a loading control. The gel is representative of two independent experiments. **D,** A time course of nanoluciferase (nLuc) synthesis in the CFPS systems prepared based on the GADD34Δ and K3L expressing (ECE) and WT HEK293T cell extracts programmed with an EMCV IRES-containing mRNA encoding nanoluciferase. All error bars represent one standard deviation of three independent replicates. (**E**) and (**F**) Representative polysome profiles of the WT HEK293T cell extract (**E**) and ECE (**F**) in the absence or presence of EMCV IRES containing nLuc mRNA template. **G,** Nanoluciferase levels from cell-free translation reactions including polyadenylated nLuc mRNAs containing different 5’ UTRs, as indicated. All templates were uncapped, except the human β-globin (*HBB*) 5’ UTR.

We describe here a highly efficient cell-free translation system based on genetically modified human HEK293T cell extracts. We find that overexpression of a truncated version of GADD34 and K3L in HEK293T cells efficiently reduces the phosphorylation of eIF2α and improves the translation activity of the resulting cellular extract, including robust formation of polysomes on exogenous mRNAs. This system can be used for 5’ m^7^G-capped and IRES-containing mRNA templates, in mRNA-dependent or transcription-translation coupled reactions. We also probed the regulation of eIF2α phosphorylation, the phosphorylation status of eukaryotic elongation factor 2 (eEF2) in these extracts and identify avenues for future optimization of the CFPS system that could enable reconstitution of translation regulatory pathways for biochemical and structural studies.

## Results

### Endogenous expression of GADD34Δ and K3L increases the translational activity of HEK293T cell extracts

To develop a quick and robust method to generate translationally active human extracts, we first tested the hypothesis that endogenously expressed proteins GADD34 and K3L might increase the synthetic activity of the *in vitro* translation system. We deleted the N-terminal 240 amino acids of human GADD34, hereafter denoted GADD34Δ, as this deletion allows high levels of GADD34 expression without compromising its enzymatic activity (*23*). By employing the Sleeping Beauty transposon stable integration system (*24*), we engineered the HEK293T human cell line to express both the GADD34Δ and K3L proteins under the control of a doxycycline-inducible promoter (Fig. 1B) (See Materials and Methods). Stable integration of the expression construct minimizes expression variation between different lysates preparations. In addition, a tightly controlled inducible promoter bypasses potential toxicities due to the overexpression of these proteins. After optimization of doxycycline levels, GADD34Δ expression was detected in HEK293T cells without visible toxic effects on the cell’s growth (Fig. 1C).

Efficient *in vitro* polypeptide synthesis requires substantial energy resources (*3, 5*). To mimic the physiological environment and bypass the potential stringency of nucleotide triphosphates in the CFPS reactions, we supplied the human cellular extract with an energy-recycling system (*2*). Although the primary energy consumption of CFPS systems is aminoacyl-adenylate formation (*25*) and, therefore, the transformation of ATP to AMP and two inorganic phosphate molecules, the creatine kinase commonly used in energy regeneration systems synthesizes ATP from ADP and creatine phosphate (*26*). To overcome this mismatch, we added rabbit myokinase, which converts the AMP released after tRNA aminoacylation to ADP, the substrate of creatine kinase, by transferring the γ-phosphate from ATP to AMP (*27*). Furthermore, since several human translation factors use GTP (*28*), we added nucleotide diphosphate kinase which can maintain the steady-state level of GTP by transferring the γ-phosphate from ATP to GDP (*29*). Altogether, these three enzymes restore the concentration of ATP and GTP, which is required for efficient cell-free translation.

The engineered HEK293T cell extract with GADD34Δ and K3L (hereafter called “engineered cell extract”) was supplemented with a nanoluciferase (nLuc) mRNA, and the *in vitro* translation was carried out under conditions described in the Materials and Methods (*8*). The synthetic activity of the CFPS system was monitored by the accumulation of enzymatically active nLuc (*30*) and was found to be ∼50-fold more active than extracts from HEK293T cells lacking GADD34Δ and K3L overexpression (hereafter WT extracts) based on the translation of enzymatically active nLuc (Fig. 1D). Moreover, the new translation system remains synthetically active for much longer periods of time, possibly suggesting the high stability of the mRNA in the CFPS system, and the increased activity of translation factors due to the modified energy regeneration system (Fig. 1D). In the above reactions, the reaction conditions were optimized separately in order to maximize the efficiency of *in vitro* protein synthesis in each extract.

To further characterize the activity of the optimized system, we measured the ability of the engineered cell extract to form polysomes on exogenously added mRNA. Extracts were incubated with polyadenylated nLuc mRNA containing the EMCV IRES, and polysome formation was monitored by sedimentation analysis on 10-50% sucrose gradients. Consistent with the increase in translation activity, polysomes were detected in the engineered cell extract-based CFPS system but not in WT cell extracts (Fig. 1E, F). The polysomes present in the reactions without adding the nLuc mRNA (Fig. 1 F, blue line) likely come from residual translation of the endogenous cellular templates in the extract. Minimizing the background translation by micrococcal nuclease treatment (7) significantly decreases the synthetic capacity of the CFPS (Fig. S1). Therefore, for the experiments described here, the cellular extracts used for the preparation of the CFPS were not treated with micrococcal nuclease.

Next, we tested if the CFPS system based on the engineered cell extract can improve translation of mRNAs with different 5’ UTRs. While the EMCV IRES containing mRNA helps drive strong protein expression without a 5’ m^7^G cap structure (Fig. 1D), its large size and complicated RNA secondary structure may limit its applications. We tested the CFPS system with nLuc mRNAs using 5’-UTRs containing the cap-independent HCV IRES (*31, 32*), CrPV IRES (*33, 34*), omega leader (*35*), or synthetic SIST sequence (*36*). However, we could not identify optimized temperature and ionic conditions for any of these 5’-UTRs that improved their translation to comparable levels as the EMCV IRES (Fig. 1G). The widely used capped human β-globin (*HBB*) 5’-UTR (*37*) shows substantial translational activity in the optimized CFPS system (Fig. 1G), but not as high as with the EMCV IRES.

### The activity of engineered HEK293T cell extracts is comparable to a HeLa-based *in vitro* translation system

The expression and purification of recombinant GADD34Δ and K3L proteins add labor and cost to preparing CFPS systems (*20, 23*). Therefore, engineering human cell lines to express these recombinant proteins should reduce the cost and time required to prepare extracts for CFPS. Consequently, we compared the translational activity of a commercially available CFPS system with cell-free translation systems prepared in-house. The commercially available system based on the S3 HeLa cell extract contains ectopically purified GADD34Δ and K3L accessory proteins (Fig. 2A). The homemade reaction was based on the ECE and contained all the required supplements (see Methods). The two translation systems were set up in parallel and in the same volume to test their activity, using nLuc or GFP activity assays to monitor the translation of EMCV IRES-containing and polyadenylated mRNAs encoding these proteins. When compared to engineered cell extracts prepared as described above, the translational activity of the commercially available HeLa translation system is not substantially higher than the in-house prepared CFPS system prepared from the engineered HEK293T cells (Fig. 2B, C). Thus the overexpression of the GADD34Δ and K3L accessory proteins in human cell lines rather than subsequent addition to cell extracts effectively promotes CFPS activity to the same extent while significantly reducing preparation time and cost (*20, 23*).

**Figure 2.**
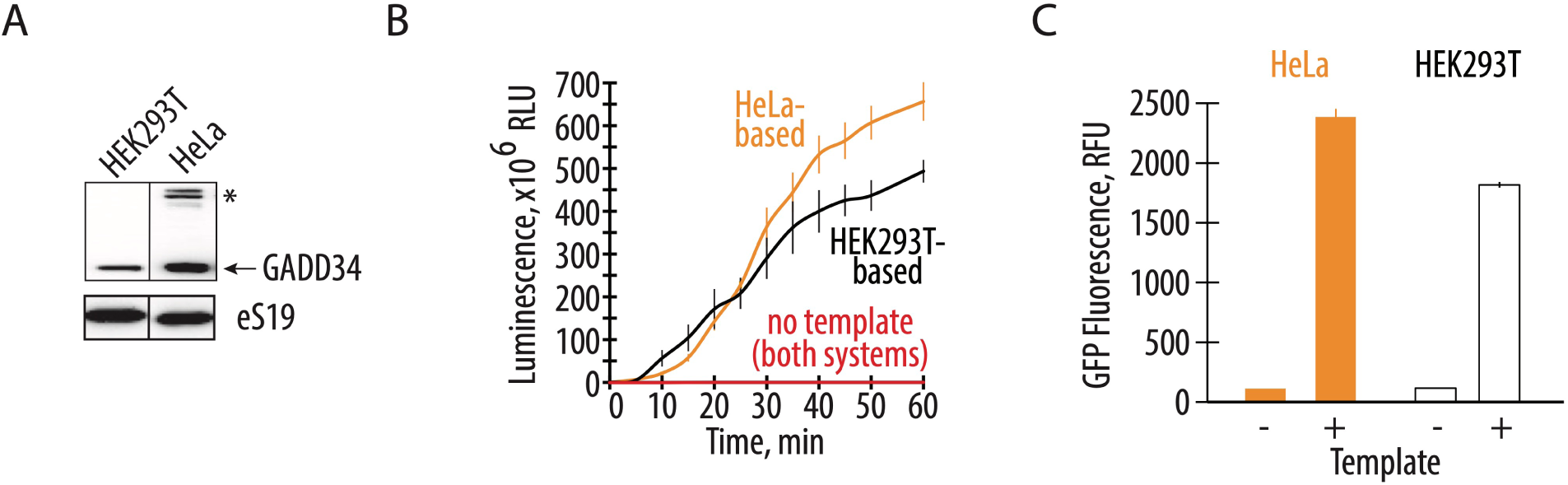
Endogenously expressed GADD34Δ and K3L increases translational activity comparable to the addition of exogenously expressed accessory proteins. **A,** Western blot showing the amount of the GADD34Δ expressed in the engineered HEK293T cells and supplemented in the HeLa-based commercial translation system. The asterisk indicates a nonspecific band in the HeLa extract. The gel is representative of two independent experiments. **B,** A time course of nanoluciferase synthesis in the CFPS systems prepared based on the engineered HEK293T cell extract and HeLa-based extract with recombinant GADD34Δ and K3L supplement. All error bars represent one standard deviation of three independent replicates. **C,** Cell-free synthesis of GFP in the two translation systems. Orange bars represent the HeLa-based extract with exogenous GADD34Δ and K3L added, while the white bars represent the engineered HEK293T cell extract with endogenously expressed GADD34Δ and K3L proteins. All error bars represent one standard deviation of three independent replicates.

### Cellular overexpression of the GADD34Δ and K3L proteins protects eIF2α from phosphorylation

The increased translational activity of the CFPS system based on the engineered cell extract seen above is consistent with the role of GADD34Δ and K3L in preventing phosphorylation of eIF2α in the *in vitro* translation reactions. To test this hypothesis, we probed the phosphorylation state of eIF2α in CFPS systems based on cell lysates from the engineered and WT HEK293T cells. By employing phos-tag gels (*38, 39*), which separate the phosphorylated and non-phosphorylated forms of eIF2α during gel electrophoresis, western blots of eIF2α revealed the baseline level of phosphorylated eIF2α in translation systems from WT and engineered cell extracts was similarly low after one hour of incubation, even though endogenous mRNAs were not removed (Fig. 3A). However, addition of exogenous mRNA to CFPS reactions significantly increased the amount of the phosphorylated eIF2α in WT extracts (up to 28%) compared to the small increase seen in engineered cell extracts containing GADD34Δ and K3L (∼5%) (Fig. 3A). This increase in WT extracts was independent of whether the 5’ UTR of the mRNA was cap-dependent (*HBB* 5’-UTR) or used the long and highly structured EMCV IRES (Fig. 3A). Remarkably, the level of phosphorylated eIF2α was highest in the translationally efficient system based on the HeLa extract supplemented with recombinant GADD34Δ and K3L proteins (∼44% and ∼51% phosphorylation without and with exogenous mRNA added, respectively) (Fig. 3B). These results clearly show that overexpression of the GADD34Δ and K3L proteins in HEK293T cells prevents eIF2α phosphorylation more efficiently compared to the ectopically added recombinantly proteins (Fig. 3A and B), even though these proteins are added at roughly the same concentration to the two CFPS systems (Fig. 2A).

**Figure 3.**
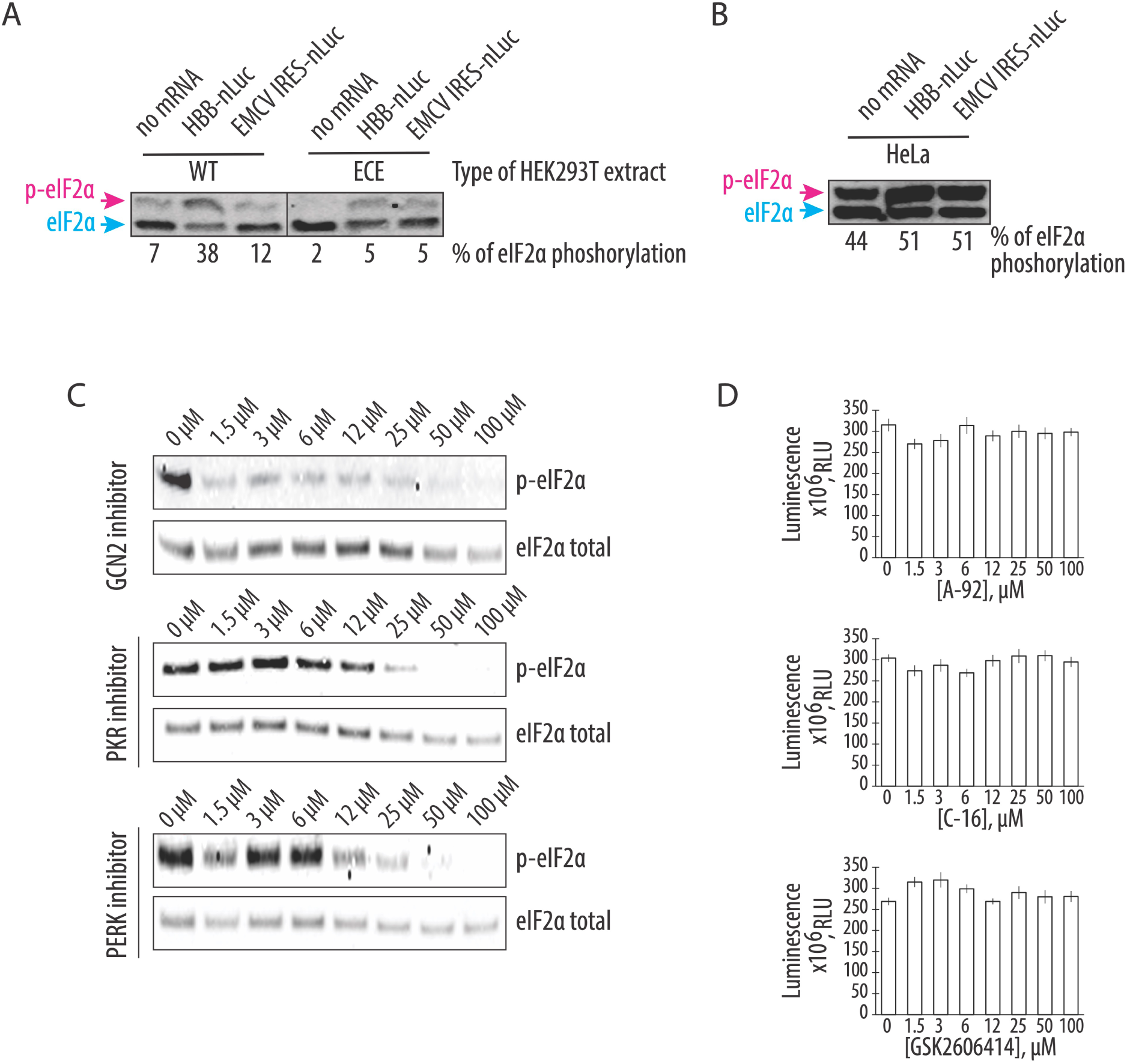
GCN2 kinase is responsible for the residual phosphorylation of eIF2α during CFPS in the engineered HEK293T extract. **A,** Western blots for phosphorylation of eIF2α in CFPS systems prepared based on the engineered and WT HEK293T cell extract. The capped human β-globin (*HBB*) and uncapped EMCV IRES-containing polyadenylated mRNAs were used to drive the synthesis of nanoluciferase. **B,** Induction of eIF2α phosphorylation in the HeLa-based commercial translation system supplemented with the same mRNAs. For both **A** and **B**, the blue arrow indicates the non-phosphorylated form of the eIF2α, while the magenta arrow indicates the phosphorylated form on the phos-tag gels. For both **A** and **B**, the gels are representative of two independent experiments. The percentage of eIF2α phosphorylation, based on the phos-tag gels, are indicated under gels (see Materials and Methods). **C,** The GCN2 but not PKR or PERK kinase inhibitor protects eIF2α from phosphorylation during CFPS. The compound A-92 (*40*) was used as a GCN2 kinase inhibitor, C-16 (*41*) for PKR kinase inhibition, and GSK2606414 (*42*) as an inhibitor of PERK kinase. The concentrations of the eIF2α-specific kinase inhibitors are indicated. Gels are representative of two independent experiments. **D,** Cell-free synthesis of nanoluciferase in different concentrations of the eIF2α-specific kinase inhibitors. All error bars represent one standard deviation of three independent replicates.

Next, we asked if the residual eIF2α phosphorylation in the CFPS system based on the engineered cell extract was mediated by a particular eIF2α-specific kinase and whether inhibition of this kinase could affect the protein synthesis activity of the system. Although the four known eIF2α-specific kinases are each activated by specific stress response signals, these signaling pathways might be dysregulated in the context of cellular extracts. For example, GCN2 kinase might be activated by the supplementation of the system with uncharged bovine total tRNA (See Materials and Methods). To test this idea, we checked the phosphorylation state of eIF2α in CFPS reactions pretreated with different eIF2α-specific kinase inhibitors (Fig. 3D). We used GCN2 kinase inhibitor A-92 (IC_50_ = 300 nM, (*40*)), PKR-specific inhibitor C-16 (IC_50_ = 210 nM, (*41*)) and PERK kinase inhibitor GSK26006414 (IC_50_ = 0.4 nM, (*42*)). Only the GCN2-specific kinase inhibitor prevented eIF2α phosphorylation at a concentration near its IC_50_ value, whereas the PERK and PKR-specific kinase inhibitors failed to prevent eIF2α phosphorylation, even at concentrations far above their IC_50_ values (Fig. 3D). While this result revealed the likely cause of eIF2α phosphorylation in the cell-free protein expression reactions, the decrease in eIF2α phosphorylation did not increase the translational activity of the engineered cell extract-based human translation system (Fig. 3D), indicating that eIF2α phosphorylation is not limiting translation in the new CFPS system.

### Additional factors may slow down translation during the elongation stage of the CFPS

Although in the ECE-based CFPS system, eIF2α phosphorylation levels do not exceed a few percent (Fig. 3A), the absence of a correlation between eIF2α phosphorylation and translational activity made us consider whether other limiting factors may affect the translation activity of the CFPS system. One potential limitation of the CFPS synthetic activity is activation of ribosome quality control (RQC) mechanisms due to defective translation in the cellular extract and collisions of translating ribosomes resulting in the formation of the disome and trisome particles (*28, 43*). To probe the formation of such particles in CFPS with the engineered cell extract, extracts were incubated with or without polyadenylated nLuc mRNA containing the EMCV IRES, digested with RNase A, and the formation of the nuclease-resistant disomes was monitored by sedimentation analysis on 10-50% sucrose gradients (Fig. 4A). Consistent with our hypothesis, translation of the residual cellular templates results in the formation of the nuclease-resistant disomes and trisomes (Fig. 4A, orange line), which may activate ribosome-dependent quality control mechanisms. Notably, translation of the nanoluciferase template does not further increase disome and trisome formation, despite the substantial increase of the polysomes (Fig. 4A, magenta and black lines, respectively). We speculate that the appearance of the disomes and trisomes due to the residual and possibly aberrant translation of the endogenous templates might be the reason for the activation of the GCN2 kinase (Fig. 3), as recently described (*44*). The presence of collided ribosomes on cellular mRNAs and subsequent activation of the ribosome quality control mechanisms might explain eIF2α phosphorylation in the engineered cell extracts, with concomitant limits on the synthetic activity of the CFPS system (Fig. 4C).

**Figure 4.**
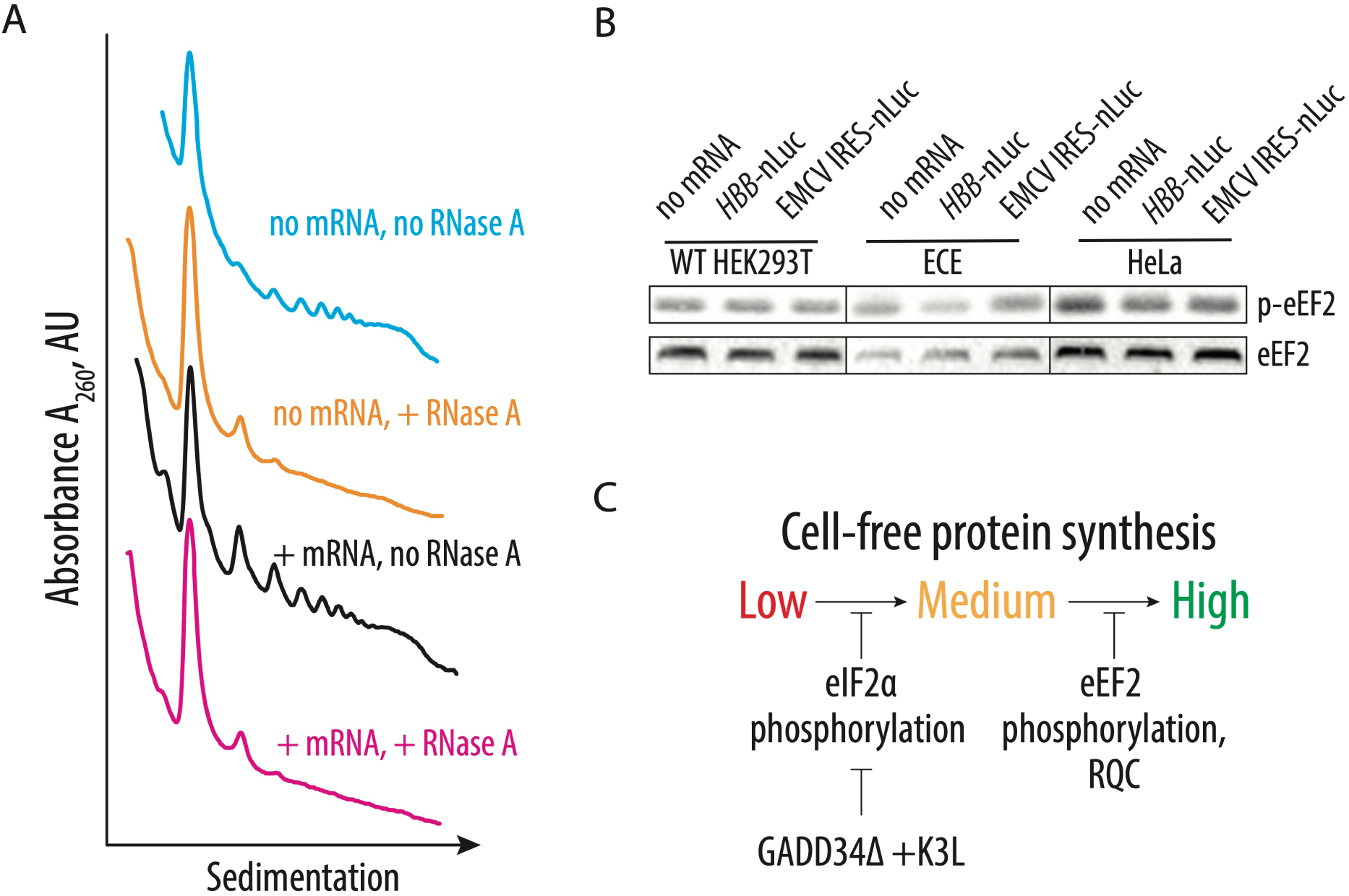
Identification of possible CFPS-limiting factors. **A,** Sucrose gradient sedimentation analysis reveals the presence of disomes and trisomes in the CFPS system based on the extract from the engineered HEK293T cell line, independent of exogenously added mRNA. **B,** Phosphorylation of eEF2 on T56 in the different translation systems. The capped human β-globin (*HBB*) and uncapped EMCV IRES-containing polyadenylated mRNAs were used to drive the synthesis of nanoluciferase. The gels are representative of two independent experiments. **C,** The proposed model for factors limiting cell-free protein synthesis in the engineered human cell extract. Phosphorylation of eIF2α inhibits translation initiation and can be bypassed by the overexpression of GADD34Δ and K3L proteins. Several factors may limit production of an even more active CFPS system by affecting translation elongation, including induction of the phosphorylation of eEF2 and activation of ribosome-associated quality control factors.

Cellular translation might be regulated not only during initiation and by RQC, but also during elongation. For example, it is possible that any degree of inhibitory phosphorylation of translation factors could potentially decrease the activity of *in vitro* translation systems. We hypothesized that CFPS might also be repressed by the phosphorylation of threonine 56 of eEF2, which prevents eEF2 binding to the ribosome and globally down-regulates translation in human cells during stress conditions (*45–47*). Elongation factor eEF2 translocates the tRNAs and mRNA through the ribosome and, therefore, is essential during each round of translation elongation. Western blot analysis with antibodies specific to the T56 phosphorylated form of eEF2 revealed increased phosphorylation of eEF2 upon *in vitro* translation in all tested CFPS extracts, independent of the presence of the GADD34Δ and K3L proteins (Fig. 4B). Although phosphorylation does not increase upon adding exogenous nanoluciferase mRNA, it may still decrease the synthetic capacity of the CFPS reactions by reducing the concentration of active eEF2 and, therefore, extending the time of each elongation cycle.

## Discussion

Here we describe the preparation of a highly-active CFPS system derived from human cells engineered to express GADD34Δ and K3L, which minimize eIF2α phosphorylation in the extract. We also implement a new energy regeneration system informed by the nucleotide pools that result from the different steps in translation. One of the most significant advantages of the translational system described here is that all components of the translational machinery are derived from human sources. While previous results in Chinese hamster ovary (CHO) cells that transiently overexpressed GADD34Δ also showed increases in the translational activity in a CFPS system (*48*), the use of CHO cells may introduce confounding variables to the study of aspects of human translation that are not conserved. Stable integration of the recombinant genes for GADD34Δ and K3L in the genome as described here should also enable more controlled preparation of cell extracts without complications due to transient transfection on cell viability (*48*). In the case of human derived CFPS systems, we find that HeLa translation systems can achieve high levels of translation (Fig. 2). Although this *in vitro* translation system is supplemented with individually purified GADD34 and K3L, the high levels of eIF2α phosphorylation seen in the HeLa extracts (Fig. 3B) indicate that HeLa translation extracts are highly dysregulated and are not suited to the study of mechanisms of translation regulation. Moreover, the composition of the commercially available translation mix is not fully defined, impeding its facile manipulation required for foundational investigations of human translation. Finally, purification of GADD34Δ and K3L from bacteria has proven to be challenging (provide citations?). Thus, the changes to lysate preparation and the translation reactions outlined here should enable the study of human translation under more physiological conditions.

The overexpression of GADD34Δ and K3L prior to cell lysis helps alleviate the inhibitory effect of eIF2α phosphorylation and supports high levels of translation exemplified by the formation of robust polysomes (Fig. 1F). Furthermore, cellular expression of these two proteins increases the reproducibility of the *in vitro* translation experiments by decreasing the initial phosphorylation of eIF2α upon lysis (Fig. 3A). Although our translational system is not treated with micrococcal nuclease to deplete endogenous mRNAs, the extract preserves the integrity of all translation machinery components resulting in high levels of translation (Fig. 2, Fig. S1), and can be used with both m^7^G-cap dependent (HBB) and IRES dependent mRNAs (EMCV IRES) (Fig. 1G). The physiological state of the eIF2α may allow the study of the effect of different factors known to influence mRNA translation, including the roles of translation initiation factors and the effects of RNA sequence and secondary structure on translation initiation (*49–57*). The fact that the present CFPS system produces polysomes in a robust manner may also allow *in vitro* studies of the mRNA circularization model of translation (*58, 59*). Finally, maintaining the physiological state of the eIF2α should also make the human-derived CFPS system valuable for assessing translational fidelity during alternative decoding events (*37*), programmed ribosomal frameshifting (*60*), and ribosome quality control pathways (*28, 61*).

The CFPS system described here provides substantial improvements to the use of *in vitro* cellular lysates for the study of human translation. However, future improvements are likely possible within this system. For example, it could be useful to find ways to deplete endogenous mRNAs from the extract, i.e. to eliminate the formation of residual disomes and trisomes (Fig. 4A) without the use of nucleases. Additionally, we observe phosphorylation of elongation factor eEF2 in all the extracts we examined (Fig. 4B). While additional biochemical analysis is required, the extracts might be further improved by decreasing eEF2 phosphorylation at residue T56 by using eEF2 kinase inhibitors (*62*). The approaches described here could be also used to prepare *in vitro* cell-free translation systems from other cell types, including immortalized cells derived from different tissues. It may also be possible to prepare CFPS systems from primary cells derived from healthy individuals as well as from patients suffering from diseases associated with translation defects, including rare ribosomopathies (*63, 64*). Efforts to reduce the number of cells needed to prepare the engineered CFPS system described here would be required but could provide unprecedented opportunities to study the molecular mechanisms underlying human translation regulation.

## Materials and Methods

### Cloning of the pSBtet-GADD34Δ/K3L expression plasmid

The plasmid backbones were prepared by SfiI (NEB) restriction of the pSBtet-Hygro vector (Addgene, 60508). The DNA fragments needed to assemble the constructs encoding GADD34Δ and K3L were PCR amplified using Q5 High-Fidelity DNA polymerase (NEB) from plasmid templates (gift of A. Pulk). For the amplification of the GADD34Δ fragment the following primers were used:

Fwd-GADD34Δ, 5’-TACCACTTCCTACCCTCGAAAGGCCTCTGAGGCCACCATGAAA GGAGCCAGGAAGACCTCCGTGT-3’

Rev-GADD34Δ, 5’-GCCAGGATTCTCCTCGACGTCACCAGCCTGCTTCAGCA GGCTGAAGTTAGTAGCGCCACGCCTCCCACTGAGGCTCAGG 3’.

For the amplification of the K3L fragment, the following primers were used:

Fwd-K3L, 5’-TGGTGACGTCGAGGAGAATCCTGGCCCCTTAGCCTTTT GTTATTCGCTCCCTA-3’

Rev-K3L, 5’-CATGTCTATCGATGGAAGCTTGGCCTGACAGGCCTCACT GGTGGCGACACATACGTTTATAA-3’.

The pSBtet-GADD34Δ/K3L expression plasmid was constructed by Gibson assembly (*65*) using the two PCR fragments and the vector backbone. The final sequence was verified by the full plasmid sequencing and the plasmid made available through Addgene (#196136).

### Cell lines and culture conditions

HEK293T cells were grown in Dulbecco’s modified eagle medium (DMEM, Gibco) in 10% FBS (Gibco), Penicillin (100 u/ml) (Gibco), and Streptomycin (100 µg/ml) (Gibco). HEK293T cells with the pSBtet-GADD34Δ/K3L construct inserted in the genome (ECE HEK293T cells) were grown in DMEM media (Gibco) supplemented with 10% Tet-system approved FBS (Gibco), Penicillin (100 u/ml) (Gibco), Streptomycin (100 µg/ml) (Gibco), and 2 mM GlutaMax (Gibco). Cells were grown at 37 °C in 5% carbon dioxide and 100% humidity. Induction of GADD34Δ and K3L expression prior to cell extract preparation is described below.

### Generation of the HEK293T cell line with a stably integrated GADD34 Δ1-240/K3L construct

A stable cell line for endogenous expression of GADD34 Δ1-240 (GADD34Δ) and K3L was generated following the procedure described previously (*24*). Using Amaxa SF Cell Line 4D-Nucleofector X Kit (Lonza), WT HEK293T cells were transfected with a transposase-encoding plasmid (pSB100X), and the pSBtet-GADD34Δ/K3L construct (see above). After two days of recovery, nucleofected cells were selected with 300 µg/ml hygromycin (Thermo Scientific). A total of three rounds of selection were performed, each with approximately three doublings. Visual observation of the cell culture monitored the efficiency of the selection. After completion of the selection, cells were regularly maintained with 150 µg/ml hygromycin.

### *In vitro* transcription reactions

*In vitro* transcription reactions were performed using PCR products generated with primers encoding a flanking T7 RNA polymerase promoter and a poly-A tail. Reactions were set up, as previously described (*66*), with 20 mM Tris-HCl pH 7.5, 35 mM MgCl_2_, 2 mM spermidine, 10 mM DTT, 1 u/ml pyrophosphatase (Sigma), 7.5 mM of each NTP, 0.2 u/ml RiboLock RNase Inhibitor (ThermoFisher), 0.1 mg/ml T7 RNA polymerase and 40 ng/μl PCR-generated DNA. After 3 h incubation at 37 °C, 0.1 u/μl DNase I (Promega) was added to the reactions, which were incubated at 37 °C for 30 min to remove the template DNA. RNA was precipitated for 2–3 h at −20 °C after adding 0.5x volume of 7.5 M LiCl/50 mM EDTA, and the resulting pellet was washed with cold 70% ethanol and dissolved with RNase-free water. The mRNA was further purified by using a Zymo RNA Clean and Concentrator (Zymo Research) before use in *in vitro* translation reactions.

DNA templates were amplified from a plasmid containing the corresponding 5’ UTR and the NanoLuc Luciferase coding sequence. Primers used for this amplification added a 30T sequence at the 3′ end to form a poly(A) tail after transcription. The *HBB* 5’ UTR containing mRNA was then capped using Vaccinia Capping enzyme (New England Biolabs) and 2′O-methylated using Vaccinia 2′O Methyltransferase (New England Biolabs). The IRES-containing mRNAs were uncapped and polyadenylated.

### Preparation of cell extracts for CFPS reactions

WT HEK293T cell extract was made as described previously (*67*). Briefly, adherent WT HEK293T cells at less than 40% confluence were collected by scraping, washed with ice-cold DPBS (Gibco), and suspended with an equal volume of lysis buffer (10 mM HEPES pH 7.4, 10 mM KOAc, 0.5 mM Mg(OAc)_2_, 5 mM DTT and 1x Protease inhibitor (Roche)). After incubation on ice for 45 min, the cells were lysed by pushing through a 1-ml syringe with a 26G needle about 15 times, followed by centrifugation at 12000g for 2 min. After centrifugation, the supernatant was aliquoted to avoid freeze-thaw cycles and flash-frozen in liquid nitrogen.

The ECE-based CFPS system was prepared according to the following procedure. First, approximately 1.2-1.5 million ECE HEK293T cells were seeded per 150 mm plate in ECE HEK293T cell-specific media, described above. The next day, the expression of GADD34Δ and K3L was induced by adding 1 µg/ml of doxycycline (Takara). After one additional day, cells were collected by scraping, washed with ice-cold DPBS (Gibco), and suspended with an equal volume of lysis buffer (10 mM HEPES pH 7.4, 10 mM KOAc, 0.5 mM Mg(OAc)_2_, and 5 mM DTT). After incubation on ice for 45 min, cells were lysed by pushing through a 1-ml syringe with a 26G needle about 15 times, followed by centrifugation at 15000g for 1 min. After centrifugation, the supernatant was aliquoted to avoid freeze-thaw cycles and flash-frozen in liquid nitrogen.

### *In vitro* translation reactions

The HeLa extract cell-free translation system was obtained from Thermo Fisher Scientific (catalog #88882) and used according to the manufacturer’s instructions. In the case of WT or ECE HEK293T reactions, the optimal concentration of the magnesium and potassium ions was determined for each new preparation of cellular extract, with the conditions below representative of typical final conditions.

For a 10 μl reaction using WT HEK293T cell extracts, 5 μl of cell extract was used in a buffer containing final concentrations of 20 mM HEPES pH 7.4, 120 mM KOAc, 2.5 mM Mg(OAc)_2_, 1 mM each of ATP and GTP, 2 mM creatine phosphate (Roche), 10 µg/ml creatine kinase (Roche), 0.21 mM spermidine, 0.6 mM putrescine, 2 mM TCEP (tris(2-carboxyethyl) phosphine), 10 μM amino acids mixture (Promega), 1 u/μl RiboLock RNase inhibitor (Thermo Scientific) and 200 ng mRNA. Translation reactions were incubated for 23 min at 30 °C and nanoluciferase activity was monitored using the Nano-Glo Luciferase Assay Kit (Promega) in a Microplate Luminometer (Veritas). To account for batch-to-batch variability, all assays presented in a given figure were carried out using the same *in vitro* translation extract and same preparation of mRNAs.

Translation reactions with the ECE-based CFPS system were set up according to a previously published procedure (*68*) with modifications. For a 10 μl mRNA-dependent reaction, 5 μl of cell extract was used in a buffer containing final concentrations of 52 mM HEPES pH 7.4 (Takara), 35 mM KGlu (Sigma), 1.75 mM Mg(OAc)_2_ (Invitrogen), 0.55 mM spermidine (Sigma), 1.5% Glycerol (Fisher Scientific), 0.7 mM putrescine (Sigma), 5 mM DTT (Thermo Scientific), 1.25 mM ATP (Thermo Fisher Scientific), 0.12 mM GTP (Thermo Fisher Scientific), 100 µM L-Arg; 67 µM each of L-Gln, L-Ile, L-Leu, L-Lys, L-Thr, L-Val; 33 µM each of L-Ala, L-Asp, L-Asn, L-Glu, Gly, L-His, L-Phe, L-Pro, L-Ser, L-Tyr; 17 µM each of L-Cys, L-Met; 8 µM L-Trp, 20 mM creatine phosphate (Roche), 60 µg/ml creatine kinase (Roche), 4.65 µg/ml myokinase (Sigma), 0.48 µg/ml nucleoside-diphosphate kinase (Sigma), 0.3 u/ml inorganic pyrophosphatase (Thermo Fisher Scientific), 100 µg/ml total calf tRNA (Sigma), 0.8 u/μl RiboLock RNase inhibitor (Thermo Scientific) and 1000 ng mRNA.

For the 10 μl coupled transcription-translation reactions, 5 μl of cell extract was used in a buffer containing final concentrations of 52 mM HEPES pH 7.4 (Takara), 35 mM KGlu (Sigma), 4 mM Mg(OAc)_2_ (Invitrogen), 0.55 mM spermidine (Sigma), 1% Glycerol (Fisher Scientific), 0.7 mM putrescine (Sigma), 5 mM DTT (Thermo Scientific), 1.25 mM ATP (Thermo Fisher Scientific), 0.83 mM each of UTP, CTP and GTP (Thermo Fisher Scientific), 100 µM L-Arg; 67 µM each of L-Gln, L-Ile, L-Leu, L-Lys, L-Thr, L-Val; 33 µM each of L-Ala, L-Asp, L-Asn, L-Glu, Gly, L-His, L-Phe, L-Pro, L-Ser, L-Tyr; 17 µM each of L-Cys, L-Met; 8 µM L-Trp, 20 mM creatine phosphate (Roche), 60 µg/ml creatine kinase (Roche), 4.65 µg/ml myokinase (Sigma), 0.48 µg/ml nucleoside-diphosphate kinase (Sigma), 0.3 u/ml inorganic pyrophosphatase (Thermo Fisher Scientific), 100 µg/ml total calf tRNA (Sigma), 0.8 u/μl RiboLock Rnase inhibitor (Thermo Scientific), 11 µg/ml T7 polymerase and 100 ng mRNA.

The ECE-based translation reactions were incubated for 60 min at 32 °C, and nanoluciferase activity was monitored using the Nano-Glo Luciferase Assay Kit (Promega) in a Microplate Luminometer (Veritas). To account for batch-to-batch variability, all assays presented in a given figure were carried out using the same *in vitro* translation extract and same preparation of mRNAs.

For the *in vitro* translation of GFP experiments, coupled transcription-translation reactions based on the HeLa and ECE extracts were supplemented with 50 ng/μl pCFE1-GFP plasmid (Thermo Scientific) according to the manufacturer protocol. After three hours incubation, reactions were transferred directly into a black 384-well plate with clear bottom (Greiner). The GFP fluorescent signal was measured using a Tecan SPARK plate reader at ex/em: 482/512 nm.

For the kinase inhibition experiments, cell-free translation systems were pretreated with the indicated concentrations of A-92 (Axon MedChem), C-16 (Cayman Chemical), or GSK2606414 (Axon MedChem) inhibitors prior to initiating the translation reactions.

### Micrococcal nuclease treatment of the HEK293T human cell lysates

Micrococcal nuclease treatment of human cell lysates was carried out according to (*8*). Briefly, before setting up the CFPS reaction, ECE cell extract was supplemented with micrococcal nuclease (NEB) at a final concentration of 15 u/ml and 0.75 mM of calcium chloride (Sigma). After the incubation at the 25 °C for 7 min, the nuclease reaction was stopped by the addition of EGTA (Sigma) solution at a final concentration of 3 mM.

### Polysome profiling

Lysates containing ∼100 pmol of ribosomes (1 A_260_ = 20 pmol) were resolved through 10%–50% sucrose gradients (20 mM HEPES-KOH, pH 7.4, 100 mM KCl, 5 mM MgCl_2_, 100 µg/ml cycloheximide (Sigma), 0.5 mM DTT (Thermo Fisher Scientific)) using a Beckmann Coulter SW41 Ti rotor at 38000 rpm (239,000 xg) for 4 °C for 2.5 h. For profiles of RNase-digested samples (*44*), lysates were treated with RNase A (Thermo Fisher Scientific) at 4 mg/L for 15 min at RT and quenched by adding 200 U of RiboLock RNases Inhibitor (Thermo Fisher Scientific). RNase-treated lysates were resolved through 10%–35% sucrose gradients using a Beckmann Coulter SW41 Ti rotor at 38000 rpm (239,000 xg) for 4 °C for 2 hr. Gradients were fractionated using a BRANDEL gradient fractionator and the absorbance at 254 nm was recorded.

### Western blot analysis

Samples were boiled in Bolt LDS Sample loading buffer (Thermo Fisher Scientific) containing Bolt/NuPAGE reduction buffer (Thermo Fisher Scientific) at 70 °C for 10 min. Samples were resolved on 4%–12% Bolt Bis-Tris Plus protein gels (Thermo Fisher Scientific) using 1x Bolt MES or MOPS SDS running buffer (Thermo Fisher Scientific) containing 1x NuPAGE Antioxidant (Thermo Fisher Scientific) according to the manufacturer’s instructions. Gels were transferred to nitrocellulose membranes using a Power Blot system (Thermo Fisher Scientific) using medium-range manufacturer parameters. Membranes were blocked with 5% nonfat dry milk (Bioworld) in PBST (10 mM Tris-HCl, pH 8 (Invitrogen), 1 mM EDTA (Invitrogen), 0.1% Triton X-100 (Sigma), 150 mM sodium chloride (Sigma)) for 1 hr at room temperature (RT) with gentle rocking. Blots were washed in PBST 3 times and then incubated with the indicated primary antibodies overnight at 4 °C with gentle rocking. Blots were washed with PBST 3-4 times over 45-60 min at RT with gentle rocking, then incubated with secondary antibodies diluted in 5% milk in PBST for 1 hr with gentle rocking at RT. Membranes were washed again with PBST, 3-4 times over 45-60 min at RT, developed using SuperSignal West Pico Plus ECL substrate (Thermo Fisher Scientific) and SuperSignal West Femto Maximum Sensitivity Substrate (Thermo Fisher Scientific) if needed and imaged on an IBright CL1000 (Thermo Fisher Scientific) system. Results shown are representative of at least two independent experiments.

### Phos-tag gels

For Phos-tag gel immunoblotting (*38*), samples were resolved on homemade 12 % Bis-Tris, pH 6.8, SDS-PAGE gels containing 50 μM Phos-tag (Wako, AAL-107) and 100 μM ZnCl_2_. Samples were boiled in Bolt LDS Sample loading buffer (Thermo Fisher Scientific) containing Bolt/NuPAGE reduction buffer (Thermo Fisher Scientific) at 70 °C for 10 min. Samples were resolved using 1x Bolt MES SDS running buffer (Thermo Fisher Scientific) containing 1x NuPAGE Antioxidant (Thermo Fisher Scientific) according to the manufacturer’s instructions. Gels were run using a constant 90 V until the bromophenol blue dye reached the bottom buffer. Before transfer to the nitrocellulose membrane, gels were soaked in 1 mM EDTA for 10 min with agitation to remove the Zn^+^ ions. Gels were transferred to nitrocellulose membranes using a Power Blot system (Thermo Fisher Scientific) using medium-range manufacturer parameters. Membranes were blocked with 5% nonfat dry milk (Bioworld) in PBST (10 mM Tris-HCl, pH 8 (Invitrogen), 1 mM EDTA (Invitrogen), 0.1% Triton X-100 (Sigma), 150 mM sodium chloride (Sigma)) for 1 hr at RT with gentle rocking. Blots were washed in PBST 3 times and then incubated with indicated primary antibodies overnight at 4 °C with gentle rocking. Blots were washed with PBST 3-4 times over 45-60 min at RT with gentle rocking, then incubated with secondary antibodies diluted in 5% milk in PBST for 1 hr with gentle rocking at RT.

Membranes were washed again with PBST 3-4 times over 45-60 min at RT, developed using SuperSignal West Pico Plus ECL substrate (Thermo Fisher Scientific) and SuperSignal West Femto Maximum Sensitivity Substrate (Thermo Fisher Scientific) (if needed) and imaged on an IBright CL1000 (Thermo Fisher Scientific) system. Results shown are representative of at least two independent experiments. The percents of the phosphorylation were estimated by comparing the intensities of the bands of phosphorylated and not phosphorylated forms of factors using ImageJ software (*69*). The background in the gel images were estimated using a rectangular region with equal dimensions positioned immediately above the band corresponding to the phosphorylated form of the proteins. A similar rectangular region was also selected below the non-phosphorylated band.

### Antibodies

The following antibodies were used in this study. Antibodies for eIF2⍺ (9722S), phospho-eIF2⍺ (S51) (9721S), eEF2 (2332S) and phospho-eEF2 (T56) (2331S) were from Cell Signaling Technology. Antibodies for RPS19 (A304-002A) were purchased from Bethyl Laboratories Inc. Anti-mouse IgG-HRP (sc-525409) and GADD34 (sc-373815) were from Santa Cruz Biotechnology. Anti-rabbit ECL IgG-HRP (NA934V) was from Thermo Fisher Scientific.

## Acknowledgments

We thank the members of the J.H.D.C. laboratory for the helpful discussion, W. Li, D. De Silva for experimental suggestions and advice, A. D. Kent and E. Pledger for critical reading of the manuscript, A. Pulk for sharing plasmids encoding GADD34Δ and K3L, N.T. Ingolia for the gift of pSBtet-Hygro and pSB100X plasmids and M. Mirabelli for helping with cell culture experiments.

## Author contributions

*Nikolay A. Aleksashin* - Conceptualization, Data curation, Formal analysis, Investigation, Methodology, Project administration, Supervision, Validation, Visualization, Writing – original draft, Writing – review and editing; *Stacey Tsai-Lan Chang* - Formal analysis, Investigation, Methodology, Writing – review and editing; *Jamie H. D. Cate* - Conceptualization, Formal analysis, Funding acquisition, Investigation, Methodology, Project administration, Supervision, Writing – original draft, Writing – review and editing.

---

**Figure S1.**
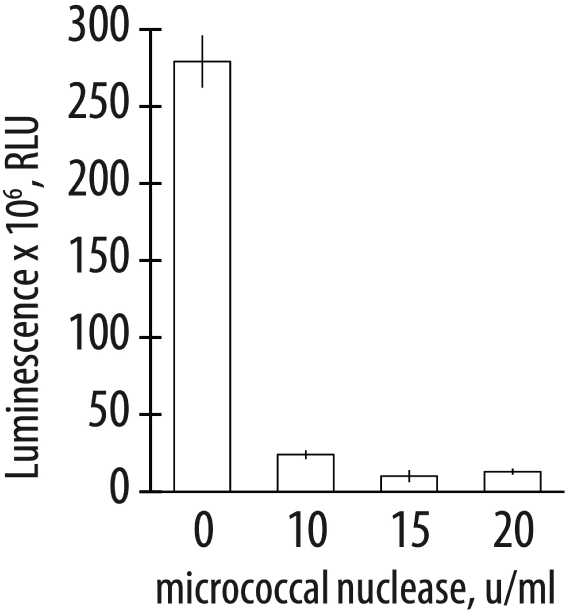
Translational activity of the human extract pretreated with micrococcal nuclease. Extracts were treated with the indicated concentrations of micrococcal nuclease prior to carrying out translation reactions with nanoluciferase mRNA. All error bars represent one standard deviation of three independent replicates.

